# Vaginal epithelial estrogen receptor α regulates vaginal pH without detectable changes in glycogen content or microbial diversity in mice

**DOI:** 10.1101/2025.10.29.685365

**Authors:** Srinivasan Mahalingam, Erin M. Carulli, Jiude Mao, Kalli K. Stephens, Jeffery A. Erickson, Wipawee Winuthayanon

**Author notes:** Corresponding author. 1030 Hitt Street, Columbia, MO 65211. **Declaration Statement:** All authors have nothing to declare.

## Abstract

Estrogen (E_2_) plays a central role in regulating the vaginal environment, but the specific contribution of epithelial estrogen receptor α (ESR1) to microbial and biochemical homeostasis has not been fully defined. In our previous work, we showed that epithelial ESR1 is indispensable for E_2_-induced epithelial proliferation and cornification, and for the suppression of MUC1 expression. Here, using mice with conditional deletion of *Esr1* in vaginal epithelial cells, called epithelial *Esr1*^d/-^, we extend these findings to demonstrate that epithelial ESR1 also regulates luminal pH without substantial changes in microbial diversity, and has no impact on total glycogen content in the vagina. Compared to control littermates, epithelial *Esr1*^d/-^ mice showed elevated vaginal pH and a subtle compositional shift in the vaginal microbiome, marked by enrichment of *Comamonadaceae* and loss of *Lactobacillus* species, without significant differences in alpha diversity. These changes parallel features of postmenopausal dysbiosis in women. Together, our findings indicate that epithelial ESR1 may modulate vaginal homeostasis by regulating vaginal pH without detectable changes in alpha diversity in mice. This work provides an insight into understanding postmenopausal vaginal dysbiosis and suggests epithelial estrogen signaling as a potential therapeutic target for genitourinary syndrome of menopause.

## Introduction

The vaginal environment is a critical determinant of women’s reproductive and gynecologic health. Disruptions in vaginal homeostasis have been associated with bacterial vaginosis, vulvovaginal atrophy, infertility, pelvic inflammatory disease, and pregnancy complications (1-4). Altered vaginal microbial communities are also linked to increased risk of sexually transmitted infections, including HIV (5). These observations underscore that mechanisms maintaining a stable microbiome and an acidic vaginal pH are central to protecting reproductive health.

In reproductive-age women, the vaginal microbiota is frequently dominated by *Lactobacillus* species, which produce lactic acid and contribute to a low vaginal pH (<4.5) that suppresses the growth of opportunistic pathogens (6-10). In contrast, dysbiosis is characterized by a reduction in *Lactobacillus*, expansion of anaerobic bacteria, and higher pH, and is strongly associated with bacterial vaginosis and reproductive tract disease (11,12). To capture bacterial community patterns, the concept of community state types (CSTs) has been developed, which classifies vaginal microbiomes into distinct clusters based on dominant taxa and diversity (13). CST frameworks highlight both the stability of *Lactobacillus*-dominated communities and the greater variability of dysbiotic states.

Estrogen (E_2_) is a central regulator of the vaginal epithelial structure and the vaginal microenvironment. In addition to regulating epithelial cell proliferation and differentiation (14,15), E_2_ has been implicated in regulating the vaginal luminal environment through several mechanisms, including glycogen accumulation and direct stimulation of proton secretion at the apical surface of vaginal epithelial cells (16-18). In addition, E_2_ and progesterone directly modulate neutrophil migration between vaginal tissues and lumen, leading to appropriate pathogen elimination (19). In states of E_2_ deficiency, such as in post-menopause, vaginal pH increases and the vaginal microbiome frequently shifts toward reduced *Lactobacillus* abundance and greater microbial diversity (18). These changes are associated with genitourinary syndrome of menopause (GSM; also known as vulvovaginal atrophy or VVA) and symptoms including vaginal dryness, irritation, and increased susceptibility to infection (20).

E_2_ responses in the vagina involve distinct functions of ESR1 in the epithelial and stromal compartments. Tissue recombination studies by Buchanan and colleagues demonstrate that stromal ESR1 mediates E_2_-induced vaginal epithelial proliferation, whereas normal epithelial stratification and cornification require ESR1 signaling in both stromal and epithelial compartments (14). Subsequent epithelial-specific deletion studies further established an intrinsic role for epithelial ESR1 in vaginal squamous differentiation (21). Consistent with these findings, we previously showed that deletion of epithelial *Esr1* disrupts vaginal epithelial differentiation, cornification, mucin 1 (MUC1) expression, and neutrophil responses (22). Although these studies establish epithelial ESR1 as an important regulator of vaginal epithelial differentiation and function, whether epithelial ESR1 also contributes to the regulation of the vaginal luminal environment, including pH and microbial composition, remains less well defined.

Here, we used an epithelial-specific *Esr1* conditional knockout mouse model called epithelial *Esr1*^d/-^ mice (22) to determine how loss of epithelial ESR1 affects the vaginal environment during estrus, a stage characterized by robust E_2_-dependent vaginal responses. This estrus model allows the contribution of epithelial ESR1 to be examined in the presence of intact ESR1 signaling in non-epithelial compartments. We hypothesized that epithelial ESR1 contributes to the regulation of the vaginal luminal environment. We assessed vaginal microbial composition and diversity, bacterial abundance, tissue glycogen content, and luminal pH. This approach allowed us to determine whether disruption of epithelial E_2_ signaling is associated with changes in the physicochemical and microbial features of the vaginal environment.

## Methods

### Mouse and Sample Collection

The work in animals was reviewed and approved by the University of Missouri ACUC, protocols #65761 and 65712. This study utilized 8-week-old mice with a selective deletion of *Esr1* in the epithelial cells of the female reproductive tract (*Wnt7a*^Cre/+^;*Esr1*^f/-^, called epithelial *Esr1*^d/-^ mice) and control littermates (*Esr1*^f/-^). Mice were housed under standard laboratory conditions with a 12-hour light/dark cycle and ad libitum access to food and water. The generation of experimental mice and genotyping were carried out as previously described (23). The deletion of epithelial *Esr*1 in epithelial *Esr1*^d/-^ mice was confirmed using an anti-ESR1 antibody (anti-ESR1 antibody, BioCare Medical, #ACA054C, RRID: AB_2651037) with a dilution of 1:400, and immunohistochemistry (IHC) analysis as previously indicated (22,24).

Mice were randomized, and the investigators were blinded to the genotype when applicable. Estrus stages were assessed using vaginal smears collected with 30 µL of sterile 0.9% saline at 08:00 - 09:00h Vaginal fluid was air-dried and stained with hematoxylin and eosin (H&E) to confirm the ovarian cycle under a light microscope.

### Vaginal glycogen assay

Vaginal tissues were collected from females (n=9-10 mice/genotype) at estrus. Tissues were weighed and immediately frozen in liquid N2 and stored at -80°C until the assay. The colorimetric glycogen assay kit II was purchased from Abcam (AB169588-1001). Glycogen content was quantified according to the manufacturer’s protocol. Briefly, 10 mg of tissue was homogenized in ddH_2_O for 10 min on ice, then boiled for an additional 10 minutes to inactivate the enzyme. Assay was performed in assay buffer, developer mix, and solution, and incubated for 30 minutes at room temperature. OD_450_ was measured using a Cytation3 microplate reader (BioTek, Winooski, VT). Glycogen content was normalized to protein content, as indicated by the BCA assay (ThermoFisher, A55864), calculated from the standard curve, and plotted as individual values with mean±SEM.

### pH measurement and vaginal swab collection

Approximately 15:00 – 16:00h of the day of estrus, vaginal pH was measured using pH strips as previously described (25). The pH paper measures pH levels from 3.0 to 5.5 with 0.5 increments (3055, Hydrion, Micro Essential Lab, Brooklyn, NY) and from 6.0 to 8.0 at 0.2 intervals (345, Hydrion). The initial pH measurement was conducted using the 6.0-8.0 range strip. If there was no color change, the 3.0-5.5 range strip was used (n=7 mice/genotype).

Subsequently, vaginal swabs were collected using sterile flock swabs (253316-H, Puritan, Guilford, ME). The tip of the cotton swab was damped with 20 µL of sterile saline, then inserted into the vaginal canal, and rotated twice clockwise and anticlockwise. Then, the tip of swaps was placed in sterile 1.5-mL Eppendorf tubes and stored at -80°C for further 16s rRNA sequencing and qPCR processing.

### Gram staining

Vaginal smears were prepared, air-dried, and heat-fixed by passing the slides (n=3 mice/genotype). Gram’s stain kit (S25344, Fisher Scientific, Waltham, MA) with Gram’s crystal violet solution for 1 minute, followed by rinsing gently with tap water. Gram’s iodine solution was applied for 1 minute, followed by rinsing. A decolorizing solution was applied dropwise until no purple color flowed from the smear. After rinsing, the smears were counterstained with safranin for 30 seconds, rinsed again, and air-dried. The image was taken using a bright-field microscope (Leica Microsystems, D1000, Deerfield, Illinois), with a 100x objective lens immersed in oil solution.

### PAS Staining

Formalin-fixed, paraffin-embedded tissue sections were rehydrated through a graded series of alcohol as previously described (n=3-4 mice/genotype) (22). Rehydrated sections were oxidized in 0.5% periodic acid (A223-25, Fisher Scientific, Fair Lawn, NJ) solution for 5 minutes, followed by rinsing in distilled water. The sections were then immersed in Schiff’s reagent (SS32-500, Fisher Scientific) for 15 minutes. After staining, the slides were washed in lukewarm tap water for 5 minutes to enhance color development and counterstained with Mayer’s hematoxylin (72711, Epredia, Kalamazoo, MI) for 1 minute. The slides were rinsed, dehydrated, and mounted for microscopic observation. The image was taken using a bright field microscope (Leica Microsystems, DMi8), with a 20x objective lens.

### Targeted V4 region 16S rRNA gene sequencing

Microbiome analyses were performed at estrus, a stage characterized by robust E_2_-dependent vaginal responses. We also attempted 16S rRNA sequencing of vaginal swabs collected at diestrus. However, four of five diestrus samples yielded insufficient sequence counts for reliable microbial community analysis. This is consistent with previous reports of lower sequencing success and reduced recoverable vaginal bacteria during diestrus compared with estrus in mice (26). Therefore, diestrus samples were not included in microbiome analysis.

Vaginal swabs were obtained from n=9 mice/genotype at the estrus stage. 16S rRNA gene sequencing was performed at the Metagenomics Center (University of Missouri). The V4 region of the 16S rRNA gene was targeted using primers U515F (5’-GTGCCAGCMGCCGCGGTAA-3’) and 806R (5’-GGACTACHVGGGTWTCTAAT-3’).

Sequencing was conducted to a depth of 100,000 reads per sample using paired-end, unique dual indexing. The data were utilized to characterize the vaginal microbial community. Data generated through the 16S rRNA gene sequencing was processed with QIIME to remove primer and index sequences. Operational Taxonomic Units (OTU) were picked with the DADA2 (27).

These OTUs were aligned to the SILVA (v138.1) reference database (28) and classified using the Naive Bayes classifier trained specifically for the V4 region, followed by constructing the feature table and performing the taxonomic classification (Qiime2-24.5). To visualize the phylum and genus, the EZBioCloud (29) web browser application was used. Fastq.gz files were directly uploaded to EZBioCloud and analyzed using the 16S-based microbiome taxonomic profiling feature. Comparative Analyzer for MTP sets was used to compare similarities and differences of taxonomic compositions between *Esr1*^f/-^ and epithelial *Esr1*^d/-^ samples.

### Microbial Classification

Vaginal microbial communities were classified using the VALENCIA (VAginaL community state typE Nearest CentroId clAssifier) algorithm, which assigns samples to community state types (CSTs) based on taxonomic composition. The analysis was performed in QIIME2 using input data with sample IDs, read counts, and formatted taxa names matching VALENCIA’s reference. The reference centroid file was used for classification (30).

To classify vaginal microbial community structures, we applied the Community State Type (CST) framework originally described by others (7). Ravel *et al*. showed that hierarchical clustering of 16S rRNA profiles from 396 asymptomatic women revealed five recurrent community state types: CST I (*Lactobacillus crispatus* dominated), CST II (*L. gasseri* dominated), CST III (*L. iners* dominated), CST IV (heterogeneous communities with low *Lactobacillus* abundance, often enriched for anaerobic bacteria such as *Gardnerella, Atopobium*, and *Prevotella*), and CST V (*L. jensenii* dominated). CST IV was further divided into subclusters reflecting different anaerobic assemblages. In our analysis, murine samples were projected against this CST reference framework to assess similarity scores and identify microbiomes from each genotype that aligned more closely with specific CST categories.

### Analysis of publicly available transcriptomic datasets

To further test our hypothesis and to identify the potential downstream targets of E_2_ through ESR1-dependent mechanism, we accessed two publicly available transcriptomic datasets of vaginal tissues from: 1) Caine et al.’s bulk RNA-sequencing analysis (GSE122591) from adult C57B6/J ovariectomized (OVX) mice after the treatment of pellet-releasing vehicle or 0.5 mg of 17β-estradiol per day for 21 days (n=5 samples/treatment) (31) and 2) Miyagawa et al.’s microarray data of control females or female lacking ESR1 in epithelial cells (*Krt5*^Cre/+^;*Esr1*^f/-^; called K5-*Esr1*KO) treated with 0.1 mg of 17β-estradiol per day for 3 consecutive days (n=2 samples/treatment) (21). For Caine et al.’s dataset, raw gene-level counts from vehicle- and 17β-estradiol-treated samples were used. Genes with fewer than 10 total reads across the ten samples were excluded. Differential expression was evaluated using DESeq2 (32) with treatment condition as the design factor and vehicle as the reference group. Microarray data were pre-analyzed and directly obtained from Miyagawa et al.’s study. A fold change was represented on a log2 scale. Genes associated with proton transport, ion transport, and glycolysis were identified. Data were then plotted using GraphPad Prism.

### Data analysis

Alpha diversity of the microbiome per sample was measured by calculating the Shannon diversity index of individual samples using QIIME 2 and represented by box plots. Wilcoxson’s test was used to determine the statistical difference between genotypes. Vaginal pH values were plotted in GraphPad Prism using mean±S.E.M. An unpaired Student’s *t*-test was performed to compare levels of pH and glycogen between genotypes, with significance at *p* < 0.05.

### Data availability

All sequencing files are available at Dryad Digital Repository; doi:10.5061/dryad.r4xgxd2sm. Coding for alpha and beta diversities was created in R Studio following the tutorial: https://github.com/microbiome-resources/Microbiome_analysis_in-_R.

## Results

We first confirmed the efficiency of epithelial ESR1 deletion in the vagina in our mouse model. The ESR1 IHC analysis showed strong nuclear staining throughout the vaginal epithelium of *Esr1*^f/-^ control mice, whereas epithelial *Esr1*^d/-^ mice displayed a marked loss of epithelial ESR1 staining, while the expression of ESR1 in stromal and other cell types remained intact (Fig. 1A). Vaginal smears from *Esr1*^f/-^ controls showed abundant adherent Gram-positive bacteria, whereas epithelial *Esr1*^d/-^ mice displayed visibly reduced Gram-positive staining and fewer bacteria attached to epithelial cells (Fig. 1B).

**Figure 1.**
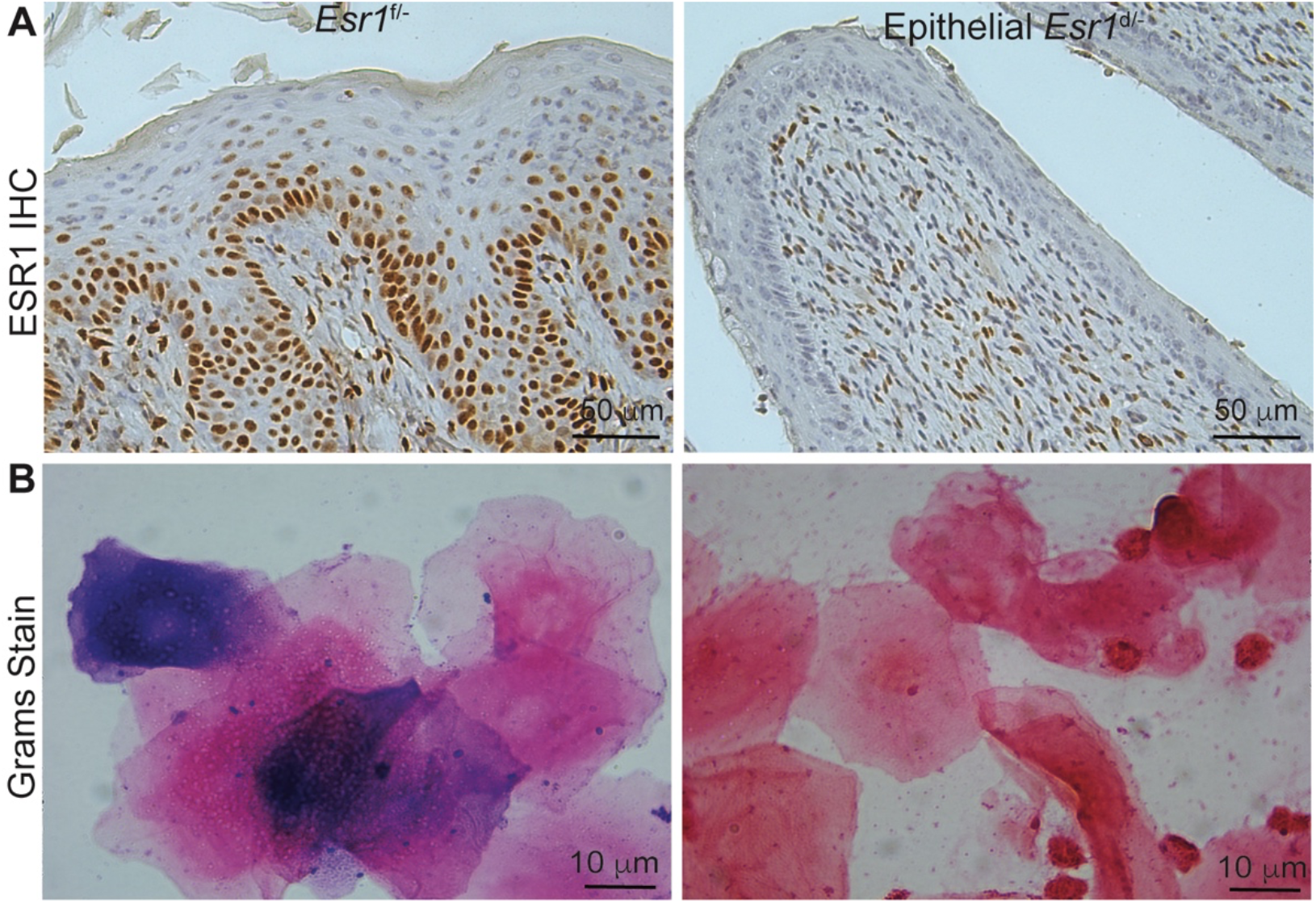
Protein expression of ESR1 in vaginal tissues and Gram stain of vaginal lavage. A. Immunohistochemistry (IHC) for ESR1 in *Esr1*^f/-^ and epithelial *Esr1*^d/-^ vaginal tissues at the estrus stage of the ovarian cycle. Scale bars, 50 µm. B. Gram staining of vaginal smears from *Esr1*^f/-^ and epithelial *Esr1*^d/-^ at estrus stage. Scale bars, 10 µm.

To characterize the impact of epithelial ESR1 on microbial community diversity, we analyzed both alpha diversity (within-sample diversity) and beta diversity (between-sample community dissimilarity) using 16S rRNA sequencing data. At the phylum level, *Esr1*^f/-^ communities were dominated by Firmicutes with contributions from Proteobacteria, while epithelial *Esr1*^d/-^ mice showed a greater representation of Actinobacteria and rare taxa (Fig. 2A-B). At the genus level, *Lactobacillus* was detected in two *Esr1*^f/-^ control samples but was not detected in epithelial *Esr1*^d/-^ samples, which showed higher relative abundances of other taxa (Fig. 2C-D).

**Figure 2.**
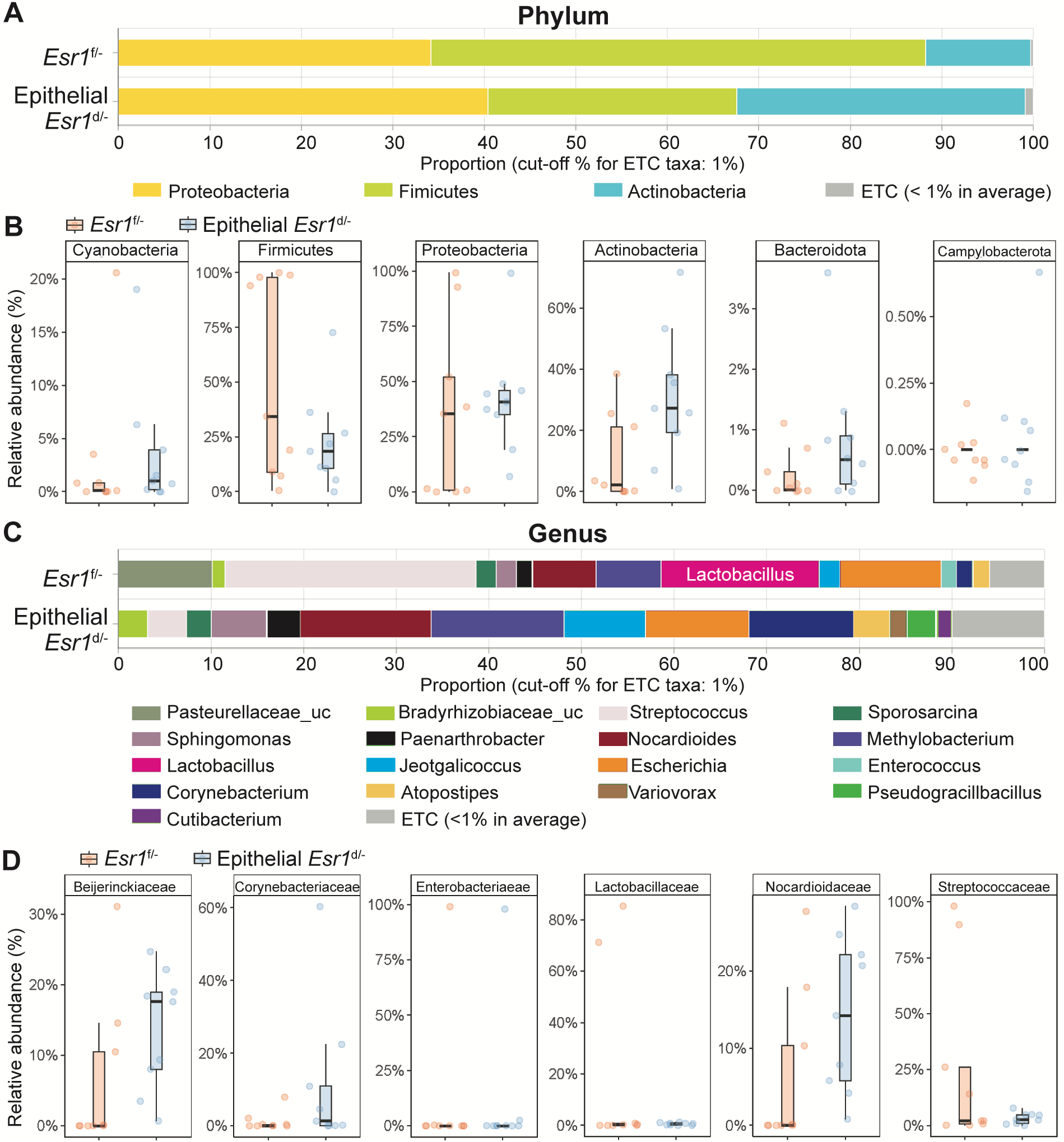
Loss of epithelial ESR1 alters vaginal microbiome composition at estrus. A. Relative abundance of major bacterial phyla (cutoff 1%) in *Esr1*^f/-^ and epithelial *Esr1*^d/-^ mice. B. The top six phyla (% relative abundance) present in *Esr1*^f/-^ and epithelial *Esr1*^d/-^ mice. C. Genus-level composition. *Lactobacillus* is detected in *Esr1*^f/-^ but markedly reduced in epithelial *Esr1*^d/-^ mice. D. The top six families (% relative abundance) present in *Esr1*^f/-^ and epithelial *Esr1*^d/-^ mice. n=9 mice/genotype. Relative abundance of each data point is indicated in B and D.

Alpha diversity reflects two properties of a microbial community: richness (the number of taxa present) and evenness (the distribution of abundances among taxa), which were assessed using observed species counts, Shannon index, Simpson index, and Pielou’s evenness. In contrast, beta diversity measures the overall compositional differences across samples, and was assessed using principal coordinate analysis (PCoA) based on Bray-Curtis dissimilarity. Alpha diversity indices (Observed, Shannon, Simpson, and Pielou’s evenness) showed no significant differences between genotypes (Fig. 3A-B), indicating that overall richness and evenness were comparable. Beta diversity analysis also showed no clear separation between genotypes despite *p* of 0.034 (Fig. 3C), suggesting that epithelial ESR1 loss may not produce a detectable difference in community structure. However, differential abundance analysis identified an enrichment of *Comamonadaceae* in epithelial *Esr1*^d/-^ mice (Fig. 3D). Samples were then classified into CSTs based on genus-level composition, following the subclusters described by Ravel *et al*. (7). CST analysis demonstrated that the vaginal microbiome in both genotypes generally aligned with CST IV subtypes (Fig. 3E), consistent with the observation that the murine vaginal microbiome is not typically dominated by *Lactobacillus*. Within CST IV subtypes, the microbiome in *Esr1*^f/-^ mice was distributed across IV-C1 and IV-C4, whereas epithelial *Esr1*^d/-^ samples were classified only as IV-C1 and had lower similarity to the IV-C1 centroid.

**Figure 3.**
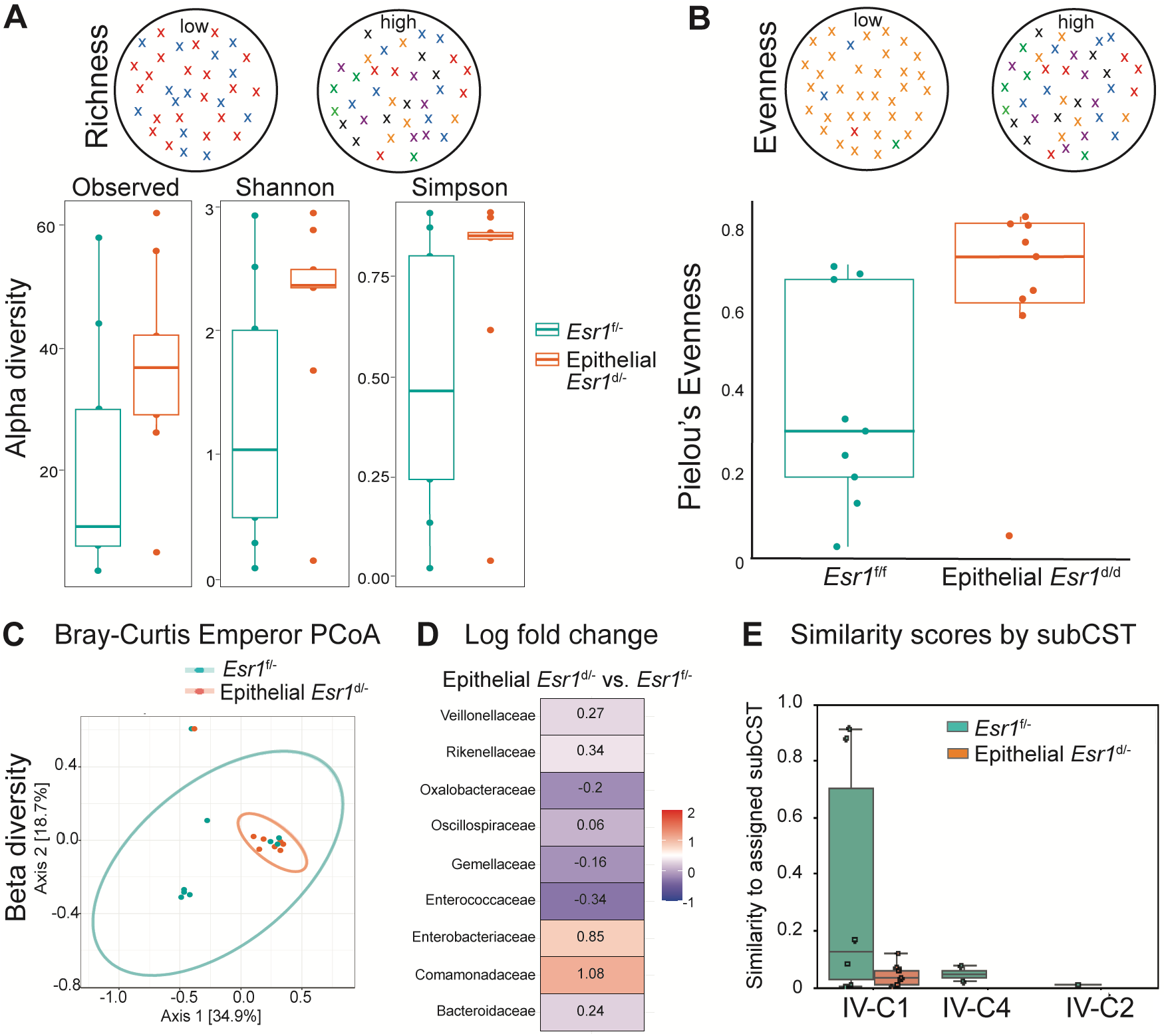
Epithelial ESR1 constrains vaginal microbiome diversity at estrus. A. Alpha diversity indices (Observed, Shannon, Simpson) in *Esr1*^f/-^ and epithelial *Esr1*^d/-^ mice. B. Pielou’s evenness in *Esr1*^f/-^ and epithelial *Esr1*^d/-^ mice. C. Beta diversity analysis using Bray-Curtis Emperor principal coordinate analysis (PCoA) in *Esr1*^f/-^ and epithelial *Esr1*^d/-^ mice. D. Log fold change of the top differentiated families between epithelial *Esr1*^d/-^ compared to *Esr1*^f/-^ mice. E. Similarity score by Community State Type IV subtyping. Samples were assigned to IV-C1, IV-C4, IV-C2, and plotted by similarity to the assigned subCST. Each data point is one mouse.

As glycogen in vaginal tissues has been proposed to influence the vaginal microbial environment (33), we examined whether loss of epithelial ESR1 altered vaginal glycogen content. PAS staining showed that PAS-positive material was present in both *Esr1*^f/-^ and epithelial *Esr1*^d/-^ tissues (Fig. 4A). Although the distribution of PAS-positive material appeared different between genotypes, quantitative measurement of total vaginal glycogen showed no significant difference between *Esr1*^f/-^ and epithelial *Esr1*^d/-^ mice at estrus (Fig. 4B). In contrast, vaginal pH was significantly elevated in epithelial *Esr1*^d/-^ mice compared to *Esr1*^f/-^ controls (Fig. 4C). These findings indicate that loss of epithelial ESR1 increases vaginal pH without a detectable change in total vaginal glycogen content.

**Figure 4.**
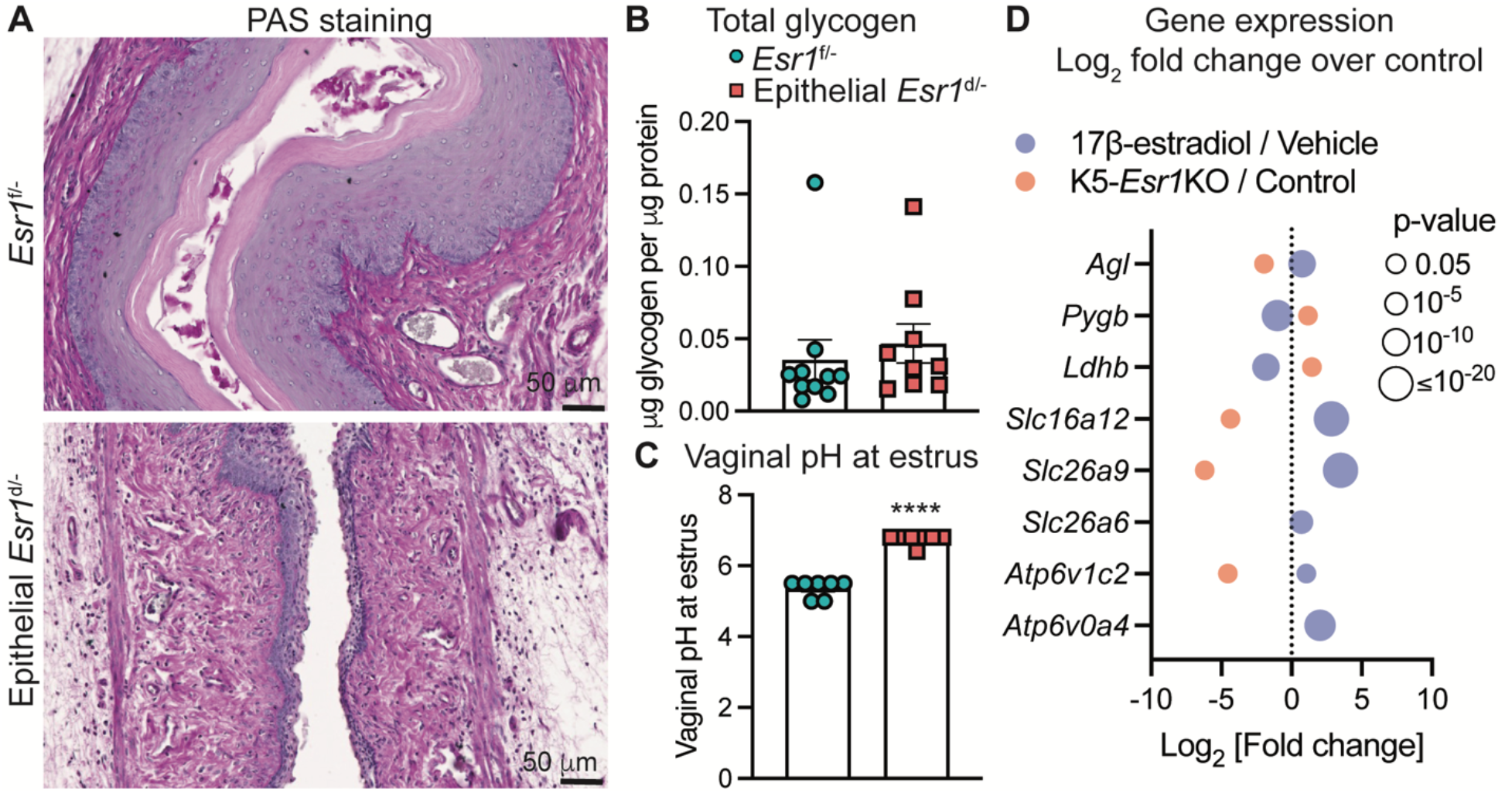
Epithelial ESR1 regulates vaginal glycogen and pH. A. Periodic Acid–Schiff (PAS) staining highlights glycogen deposition (bright purple) in *Esr1*^f/-^ and epithelial *Esr1*^d/-^ vaginal tissues at the estrus stage of the ovarian cycle. n=3-4 mice/genotype. Scale bars, 50 µm. B. Total glycogen content per protein content within vaginal tissues collected from *Esr1*^f/-^ and epithelial *Esr1*^d/-^ females at estrus. n=9-10 mice/genotype. C. Vaginal pH measurements in *Esr1*^f/-^ and epithelial *Esr1*^d/-^ mice collected at estrus. n=7 mice/genotype. Bars represent mean ± SEM; individual animals shown as points. \*\*\*\**p* < 0.0001, unpaired Student’s *t*-test. D. 17β-estradiol/epithelial ESR1-associated expression of genes involved in epithelial ion transport and carbohydrate metabolism in mouse vaginal tissue using publicly available vaginal transcriptomic datasets from Caine et al. (2019) and Miyagawa et al. (2015). Fold changes were shown as a Log2 scale for 17β-estradiol-vs. vehicle-treated vaginal tissue (blue) and K5-*Esr1*KO knockout vs. control vaginal tissue (red). Positive values indicate increased expression and negative values indicate decreased expression relative to the respective control group. Bubble size represents the nominal *p*-value. Genes not represented in the Miyagawa et al. dataset (*Slc26a6* and *Atp6v0a4*) are not shown for that comparison.

To determine potential roles of E_2_ and ESR1 in vaginal epithelial cells on the regulation of pH and glycogen, we next evaluated two different publicly available datasets: pharmacological treatment by Caine *et al*. 2019 (wild-type vaginal tissues after OVX and treated with 17β-estradiol or vehicle) and genetic deletion model by Miyagawa *et al*. 2015 (vaginal tissues from K5-*Esr1*KO or control treated with 17β-estradiol). These genes are involved in ATPase H+ transporting V0 and V1 (*Atp6v0a4* and *Atp6v1c2)*, bicarbonate and chloride transport proteins (*Slc26a6, Slc26a9*, and *Slc16a12*), lactate dehydrogenase (*Ldhb*), and glycogen breakdown (glycogen phosphorylase; *Pygb* and amylo-alpha-1,6-glucosidase, 4-alpha-glucanotransferase; *Agl*). Overall, the treatment of 17β-estradiol appeared to increase the expression of genes involved in acidification, while those involved in glycogen breakdown, with the exception of *Agl* (Fig. 4D). However, in the absence of ESR1 in vaginal epithelial cells, the expression profile of these genes was the opposite of those in 17β-estradiol-treated vaginal tissues. These data suggest that ESR1 was involved in the regulation of genes important in ion channels modulating pH and glycogen content in the vaginal. However, how subtle changes in bacterial composition in the epithelial *Esr1*^d/-^ mice contribute to increased vaginal pH remains unclear.

## Discussion

The vaginal epithelium is a hormonally responsive barrier that contributes to the local physicochemical and microbial environment of the female reproductive tract. E_2_-dependent changes in epithelial differentiation, glycogen availability, and acidification have been proposed to influence vaginal microbial communities and help maintain a low vaginal pH. Here, we investigated the specific contribution of epithelial ESR1 to these features of the vaginal environment during estrus. Loss of epithelial ESR1 increased vaginal pH without a detectable difference in total vaginal glycogen content. Alpha diversity indices also did not differ significantly between genotypes, although several taxonomic differences were observed. Beta diversity did not show clear separation between genotypes. Thus, the present data do not establish that pH regulation is independent of glycogen metabolism or microbial composition.

Glycogen availability has long been proposed as an important link between E_2_ signaling, the vaginal microbiota, and luminal pH. Glycogen breakdown products can serve as substrates for lactic acid production by vaginal commensal bacteria (34). Epithelial ESR1 regulates glucose transport (35) and may drive glycogen accumulation and influence stromal metabolism through paracrine signaling (23). Although PAS staining in our study suggested slight differences in the distribution of glycogen within the vaginal tissue, quantitative analysis showed no significant difference in total glycogen content between genotypes. Thus, total glycogen abundance alone does not explain the elevated vaginal pH observed in epithelial *Esr1*^d/-^ mice. E_2_ may also regulate vaginal pH through glycogen-independent mechanisms. In human vaginal-ectocervical epithelial cells, E_2_ directly stimulates proton secretion through apical V-ATPase, providing a mechanism by which epithelial E_2_ signaling can contribute directly to luminal acidification (16,17).

Candidate-gene expression patterns from pharmacological and genetic mouse studies are consistent with a possible role for epithelial ion transport in vaginal acidification. However, transcript abundance does not establish V-ATPase assembly, apical localization, or functional proton secretion (21,31).

Although vaginal health in women has often been equated with *Lactobacillus* dominance (36,37), emerging evidence suggests that this paradigm does not fully capture the diversity of normal states. In large cohort studies, up to one-third of asymptomatic reproductive-age women harbor communities with lower *Lactobacillus* abundance and higher proportions of anaerobes, yet remain clinically healthy (7,38). Ethnicity, host genetics, and sociodemographic factors contribute to this variation (38), and not all diverse communities confer the same risk. Thus, while *Lactobacillus* dominance supports a low pH environment and is protective in many women, the absence of *Lactobacillus* alone is not a universal marker of disease. These considerations are especially important when interpreting mouse studies, since the murine vaginal microbiome is rarely *Lactobacillus*-dominated and instead comprises diverse facultative and obligate anaerobes (25). In our study, a high relative abundance of *Lactobacillus* was observed in two of nine control samples, whereas its average abundance in epithelial *Esr1*^d/-^ samples was below the 1% plotting threshold. Therefore, the findings indicate a marked reduction rather than a confirmed complete absence of *Lactobacillus* in the epithelial *Esr1*^d/-^ mice.

The higher relative abundance of *Comamonadaceae* in epithelial *Esr1*^d/-^ mice, with no significant changes in alpha diversity, suggests that epithelial ESR1 may primarily influence microbial composition rather than overall diversity. Members of this family are aerobic or facultatively anaerobic chemoheterotrophs that proliferate in neutral-to-mildly alkaline environments (39,40). Their increased abundance in epithelial *Esr1*^d/-^ mice may reflect the elevated vaginal pH observed in this model.

Although many studies using OVX models have established that systemic E_2_ deficiency leads to loss of epithelial glycogen, increased vaginal pH, and microbial shifts toward higher diversity, these models confound effects on stroma, systemic hormones, and immune compartments. For example, OVX frequently induces vaginal atrophy, thinning of the epithelium, and broad endocrine changes (41-43). Epithelial-specific deletion of *Esr1* reproduced the elevated pH component of the OVX phenotype without systemic hormone depletion. Unlike many OVX models, however, epithelial *Esr1* deletion was not accompanied by a detectable reduction in total vaginal glycogen or a significant difference in alpha diversity. These findings identify epithelial ESR1 as a contributor to vaginal pH regulation but do not establish that all E_2_-dependent effects on glycogen or the microbiome are mediated through epithelial ESR1.

Several limitations should be acknowledged. Our experiments were performed in a single mouse strain, and mouse vaginal microbiota differ from those of women, limiting direct taxonomic translation. Future studies using human vaginal epithelial organoids, vagina-on-a-chip models, or humanized mouse models will help test whether similar mechanisms operate in women. In addition, integrating functional assays of lactate metabolism, epithelial transport, and immune signaling will further clarify how ESR1 orchestrates the vaginal ecosystem.

In summary, loss of epithelial ESR1 increased vaginal pH at estrus without a detectable difference in total vaginal glycogen content or alpha diversity. Selected taxonomic differences and candidate transcriptomic patterns suggest possible effects on microbial composition and epithelial ion transport, but functional studies are required to establish the responsible mechanisms and direction of causality. These findings extend the established role of epithelial ESR1 in vaginal differentiation and barrier integrity and identify epithelial ESR1 as a potential contributor to the regulation of the vaginal luminal environment.

## Acknowledgements

This study was supported by the Eunice Kennedy Shriver National Institute of Child Health & Human Development (NIH/NICHD) award numbers R01HD097087 and R01HD108198 to W.W.

## Author contributions

SM and WW conceptualized the research; SM, EMC, JM, KKS, JAE performed the experiments; SM and WW performed data analysis; SM and WW drafted the manuscript; SM, EMC, JM, KKS, JAE, and WW edited and approved the final manuscript.

